# Feature-based gating of cortical information transmission

**DOI:** 10.1101/2021.10.08.463709

**Authors:** Sonia Baloni Ray, Daniel Kaping, Stefan Treue

**Author notes:** Author contributions: ST, DK and SBR designed the study; SBR and DK recorded data and performed analysis; ST, SBR and DK interpreted the data and wrote the publication. These authors contributed equally (Sonia Baloni Ray, Daniel Kaping).

## Abstract

In highly developed visual systems, spatial- and feature-based attentional modulation interact to prioritize relevant information and suppress irrelevant details. We investigated the specific role and integration of these two attentional mechanisms in visual cortical area MST of rhesus monkeys. We show that spatial attention acts as a gate for information processing by providing *unimpeded high-gain pass-through processing* for all sensory information from attended visual locations. Feature-based attentional enhancement does not only show the known dependency on a match between the attended feature and a given cell’s selectivity, but surprisingly is *restricted to those features for which a given cell contributes to perception*. This necessitates a refinement of the feature-similarity gain model of attention and documents highly optimized attentional gating of sensory information for cortical processing. This gating is shaped by neuronal sensory preferences, behavioral relevance, and the causal link to perception of neurons that process this visual input.

## Introduction

Optimal perception and action in a complex environment depend on an internal neural representation of the sensory input. Correspondingly, sensory information processing in highly developed nervous systems, such as that of humans and other primates is characterized by powerful sensors that have evolved to provide the most comprehensive information possible about the environment. This representation not only encodes the detailed information provided by high-performance sensory systems, but should ideally also prioritize relevant information and suppress irrelevant details to optimize the use of limited processing resources. Otherwise, the torrent of incoming information would overtax even the most highly developed central processing capacity. High-performance sensory systems are therefore complemented by an equally sophisticated attentional system, employing an interaction of top-down (attentional) with bottom-up (sensory) processes and a combination of spatial- and feature-based attentional modulation to selectively allocate processing resources to the most relevant aspects of the input. This filtering and selection maximizes the gain for neurons representing stimuli that are (potentially) relevant and lowering the gain of other neurons, creating a weighted representation of the environment combining stimulus saliency (bottom-up) with the internally determined behavioral relevance of the corresponding sensory input (top-down). The emerging integrated saliency map reflects this top-down biased attention-driven stimulus selection, at the expense of a representation that solely aims to provide an accurate one-to-one copy of the sensory environment (1).

For the visual cortex of primates two main forms of attention have been identified, namely spatial attention and feature-based attention. The former serves to select and enhance stimuli and their representation based on spatial location. Correspondingly, the gain of neurons whose receptive field overlaps the spatial spotlight of attention is maximized and their spatial profile is biased towards the location and size of the attentional spotlight, optimizing the processing for an unimpeded high-gain passthrough of sensory information from relevant locations in the environment (2–11). Processing of information from receptive field locations outside the current spatial spotlight of attention on the other hand is impeded by reducing the gain of those neurons (12–14).

Feature-based attention, in contrast, affects information processing *across* the visual field. Rather than the pass-through routing of information caused by the spatial spotlight of attention the feature-based attention system up- or down-regulates the gain of neurons in sensory cortex based on the match between their feature-preferences and the currently attended feature set (feature-similarity gain model of attention, (15–17). Employing such modulation, the visual system enhances the representation (or ‘integrated saliency’) of stimuli that match the attended features (such as a particular direction of motion or a specific color) and suppresses the processing of all non-matching stimuli across the visual field, independent of where the current spotlight of attention is directed.

Separate roles and effects of spatial and feature-based attention have been suggested before (e.g. (18–21)) and they are in line with the results from a recent series of experiments that have shown that while spatial attention (i.e. attentional pointers in a retinotopic map) is rapidly shifted in line with saccadic eye movements this does not apply to feature-based attention (22–24).

Throughout early and mid-level visual cortex, neurons have well-defined spatial receptive fields, but show tuning for different non-spatial visual dimensions (such as orientation, color, direction of motion or stereoscopic disparity). This allows us to test the prediction that the ‘pass-through’ aspect of spatial attention affects neurons based on their receptive field location (but independently of any other visual preference and its behavioral relevance), while feature-based attention enhances the gain of only those neurons which feature-preference matches the organism’s current visual task at hand (independently of the neurons’ spatial preference). We further hypothesize that the preference of a sensory neuron for a given feature (as expressed in its tuning curve) is a necessary but not sufficient condition for a modulation by feature-based attention. Specifically, neurons that show tuned responses to a particular stimulus dimension without directly contributing to perceptual tasks involving that dimension might not be affected by feature-based attention.

We tested these hypotheses by recording from neurons in area MST of rhesus monkeys. MST is a mid-level extrastriate visual area where a large proportion of neurons are known to be tuned to linear and to more complex spiral motion patterns (25–28). Furthermore, MST neurons have been shown to be modulated by covert voluntary spatial attention (7,29,30). We focused on neurons showing a multiplexed stimulus selectivity, i.e. independent tuning for both linear and spiral motion stimuli. During the recordings the animals performed a task that engages both spatial and feature-based attention. Our data show a “pass-through” gating effect of spatial attention and a feature-based attentional modulation for spiral motion stimuli. Most interestingly though, feature-based attentional modulation showed a dissociation for the two stimulus types, being absent for linear motion stimuli, despite the prevalence of such effects in the preceding cortical area MT. This documents the predicted differences in spatial and feature-based attention, but additionally suggests that the linear motion tuning observed in MST is not directly contributing to linear motion perception.

## Results

Neuronal response selectivity for a particular stimulus dimension (such as orientation, direction of motion, etc.) is usually considered a necessary (but not sufficient) condition for the functional recruitment of a given neuronal population or cortical area in a perceptual task involving that stimulus dimension. Correspondingly, a multitude of studies have linked such neuronal selectivity in primate visual cortex to perceptual performance (31–33). Additionally, the magnitude of attentional modulation of neuronal activity has been shown to reflect the similarity of a neuron’s sensory preference to the currently attended location and stimulus features (the feature-similarity gain model of attention (15,34). We wanted to test if such functional recruitment of neurons and their attentional modulation also applies for neurons with multiplexed stimulus selectivity. For this, we measured responses of MSTd neurons while rhesus monkeys were engaged in a spatial or feature-based attention task. Two representative types of visual motion patterns - linear and spiral moving random dot patterns (RDPs) - were used, as MSTd neurons show simultaneous (but independent) tuning for both these stimulus types.

Prior to our measurements of attentional modulation, we determined the selectivity (preferred direction and speed) to complex spiral motion stimuli for each isolated neuron as a prerequisite for further analysis. We analyzed well-isolated single cell responses from 105 MSTd neurons with spiral motion tuning from two monkeys. Out of this population, 48 neurons additionally showed tuning to linear motion stimuli. This linear motion tuning was similar (in dynamic range and tuning width) to the same cell’s spiral motion tuning (see supplementary Figure1).

This subset of the neuronal population tuned to both spiral and linear motion stimuli offers the possibility to compare spatial and feature-based attentional modulation for the two stimulus types for a given cell. Hence, for these 48 neurons spatial and feature-based attentional tasks were recorded for both spiral and linear RDPs.

Just like the magnitude of location-bound attentional enhancements in the spatial attention condition, the magnitude of feature-based attentional modulation is subject to the similarity between the currently attended feature and the preferences of the neuron under study (15,34). The animals were instructed by a stationary cue, prior to the onset of coherent motion, to attend either to the stimulus inside the RF (always moving in the neuron’s preferred direction) or outside the RF (moving either in the preferred or anti-preferred direction), while ignoring the uncued stimulus (Figure 1). We determined the feature-based attentional response enhancement of 105 neurons (48 neurons tuned for both linear and spiral motion) by comparing the firing-rates between those trials where the target stimulus was the RDP outside the RF, moving either in the preferred or anti-preferred direction (mimicking the approach used in (15)). The average behavioral performance (mean % ± SD) of the two monkeys for the three conditions was computed. For MSTd recording sessions with spiral motion stimuli the performance was 90% ± 11 for attention inside the RF to the preferred direction, 88% ± 13 for attention outside the RF to the preferred direction and 88% ± 12 for attention outside the RF to anti-preferred direction. The performances for sessions with linear motion in MSTd and MT are provided in supplementary table. The feature-based attentional response modulation of an example neuron to linear and spiral motion stimuli is shown in figure 1. Figure 1C and 1D represent single unit responses for spiral and linear RDPs respectively. Spiral motion responses of this neuron were higher when the attended motion outside the receptive field moved in the preferred direction as opposed to the condition when the anti-preferred direction was attended (Fig 1C), documenting a feature-based attentional modulation of spiral motion processing in line with previous finding and the feature-similarity gain model (15). A feature-based attentional modulation of linear motion (Fig 1D), on the other hand, was not apparent in this neuron.

**Figure 1:**
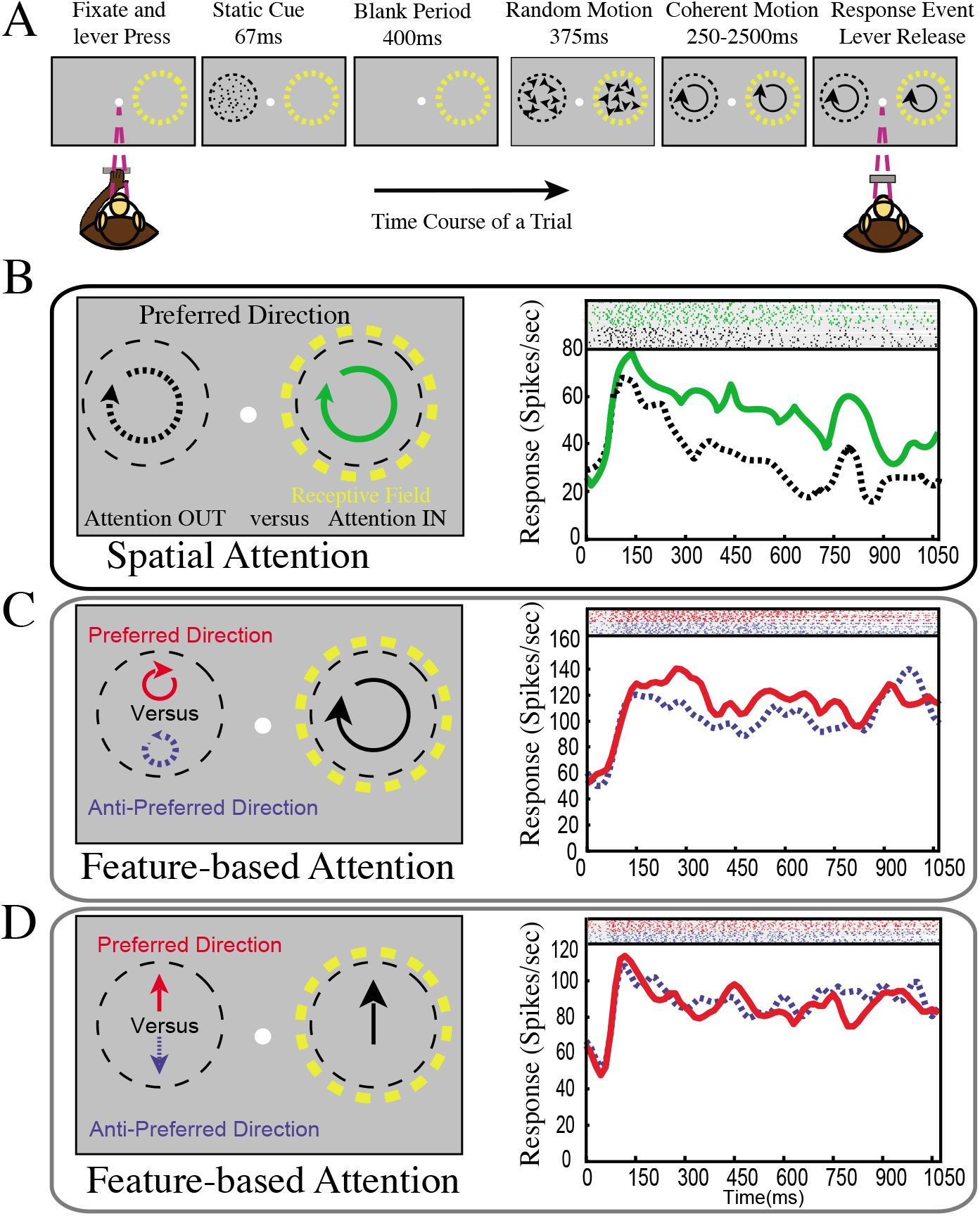
Task details. A Behavioral paradigm: The monkeys initiated each trial by directing and maintaining their gaze on a centrally presented fixation point and holding a touch sensitive lever. After trial initiation a static cue appeared for 67 ms at an eccentric location, cueing the animal to covertly shift attention towards this target location, either within the receptive field (yellow dotted circle added to figure for illustrative purposes) or in the opposite hemifield. The cue presentation was followed by a blank period for 400ms to measure baseline activity. To reduce transient motion onset responses, random motion, both inside and outside the receptive field, was presented briefly (375ms) prior to the onset of coherent motion stimuli (either spiral or linear motion). In order to obtain a liquid reward, the monkeys had to respond to a transient speed increment (250-2500ms after onset) of the target stimulus by releasing the lever, ignoring any speed changes in the distractor. B Spatial attention effects: Stimulus configuration for determining spatial attention modulation by cueing either the stimulus inside the receptive field (yellow dotted circle) moving in the preferred direction of the neuron (green circular arrow) or the stimulus outside the receptive field (black dotted circular arrow) also moving in the preferred direction. In a given trial the stimuli were either both moving in a spiral or a linear motion. The spike density function and raster plots on the right show a single unit response to the two different conditions with spiral motion stimuli. C Feature-based attention effects: Stimulus configuration for the feature-based attention condition with spiral motion. Attention was always directed to the stimulus outside the receptive field (opposite hemifield of yellow dotted circle) to either preferred direction (red) or anti-preferred direction (blue dotted). Inside the receptive field (yellow dotted circle) the stimulus always moved in the preferred direction to ensure a strong sensory response. The right panel shows an example neuron’s spike density and raster plot for responses while the target stmulus was moving either moving in the preferred (red) or anti-preferred direction (blue dotted). D Feature-based attention example for the linear motion configuration. This panel is identical to panel C, except that linear motion stimuli were presented. The right panel shows the neuronal response for the same neuron as shown in panel C, but for linear motion stimuli.

To determine the effects of feature-based attention across our population of recorded cells we quantified the attentional modulation of firing-rates computing a widely used attentional index for every neuron. The firing rates of each cell were determined for an epoch of 600ms, starting 300ms after the onset of the RDPs (gray shaded area in figure 2A and 2C). The distribution of attentional indices is shown in figure 2B (for spiral RDPs) and 2D (for linear RDPs). Mean feature-based attention index for the spiral motion attention task of 105 neurons (Figure 2B) was 0.046, corresponding to a ~10% increase in activity (p< 0.001, signed-rank test). The observed enhancement is comparable with feature-based attentional enhancements previously reported in other visual areas (e.g. MT (15) and V4 (35)).

**Figure 2:**
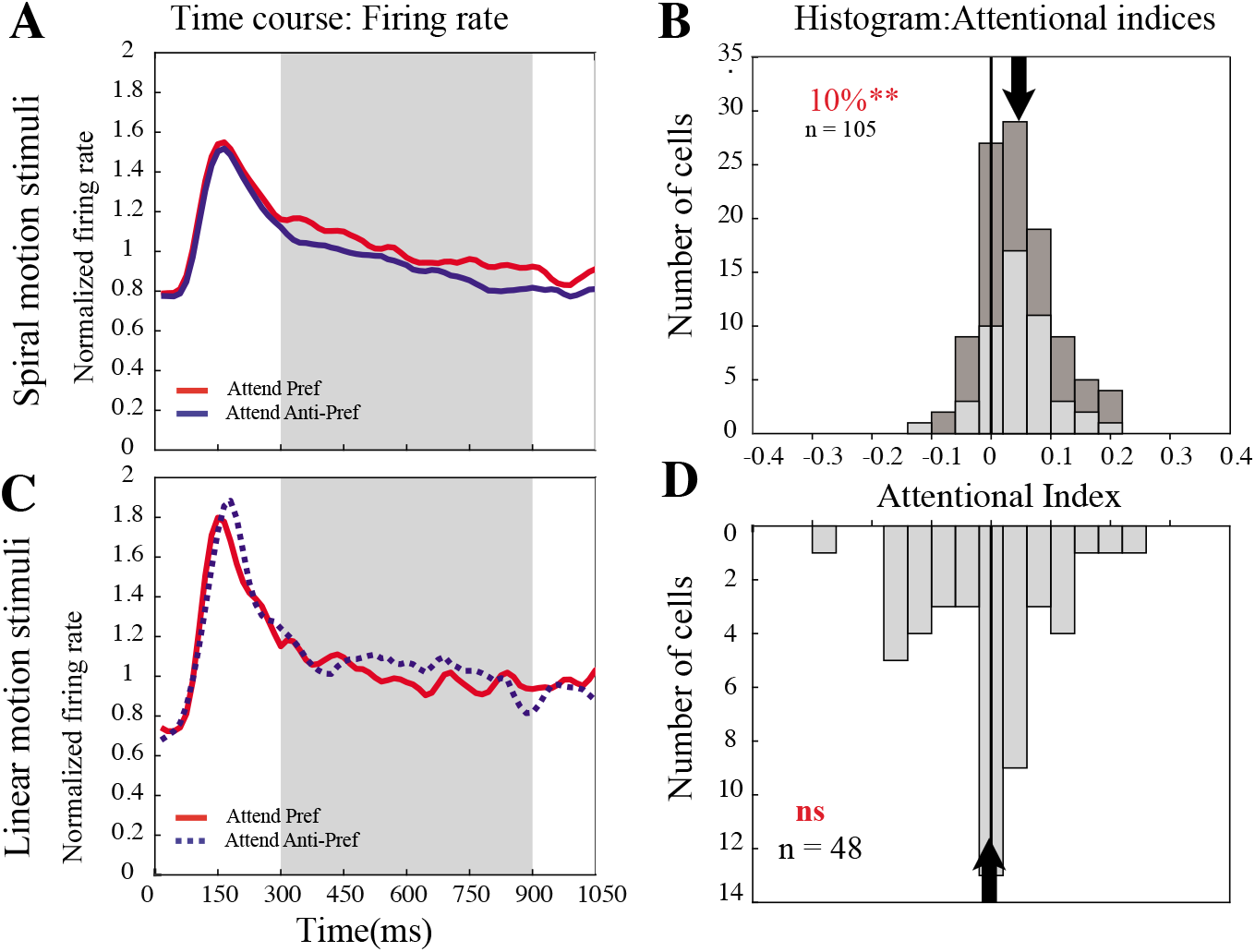
Feature-based attention effects with linear and spiral motion stimuli in MSTd. A) Population normalized spike density functions, showing the responses to a target moving in the preferred (red trace) and anti-preferred (blue dotted trace) spiral motion direction outside the receptive. Gray shaded area represents the 600 ms time period of sustained activity chosen to evaluate attentional indices, starting 300 ms of after the onset of coherent spiral motion. B) The histogram shows feature-based attentional index for spiral motion stimuli for 105 neurons recorded in macaque area MSTd for the time period represented by the gray shaded area in panel A. Binning is according to the attentional index AI = (R_pref_ − R_anti-pref_) / (R_pref_ + R_anti-pref_), where Rpref is the response corresponding to attention to preferred direction and Ranti-pref is response for attention to the anti-preferred direction. The mean attentional index for the population is 0.043 (arrow above the histogram) corresponding to an average attentional enhancement of 9% (signed-rank test, p<0.001). The light gray part of the histogram represents feature-based attentional modulation with spiral motion stimuli for the subset of 48 neurons tuned for both spiral and linear motion. C) Population normalized spike density functions, showing responses to a target moving in preferred (red traces) and anti-preferred (blue dotted trace) linear motion direction outside the receptive field. The data correspond to a subset of 48 neurons for which data were recorded for both spiral and linear motion stimuli. Gray shaded area represents the 600 ms time period of sustained activity (same time epoch as in panel A) used to compute attentional indices, starting 300 ms after the onset of coherent spiral motion. D) The histogram shows feature-based attentional index with linear motion stimuli for the subset of 48 neurons, computed for the gray shaded time period in panel B. No significant attentional modulation was observed for these 48 neurons with linear motion stimuli, even though the same neurons showed a significant (signed-rank test, p<0.01) feature-based attentional modulation (avg: 10%) with spiral motion stimuli (Light gray shaded histogram in panel B).

We next calculated attentional indices for linear motion stimuli for the sub-set of 48 MSTd neurons selective for both spiral and linear motion stimuli. Surprisingly, no systematic firing-rate enhancement for the linear motion feature-based attention (p = 0.918) condition was observed (Figure 2D). This is even though the same subset of 48 neurons shows a significant feature-based attentional enhancement of ~10% (p<0.001; figure 2B, grey shaded histogram) for spiral motion stimuli that was significantly different from that for linear motion stimuli (p = 0.0051, Wilcoxon ranksum test).

To rule out that the lack of feature-based attentional modulation for linear motion was because the animals allocated feature-based attention only to spiral motion we recorded an additional 60 neurons from area MT in the same two animals engaging in a linear motion feature-based attention task. The population spike density function and attentional index histogram (supplementary Figure 2) show a significant feature-based attentional modulation of ~6% (p = 0.0055) in area MT for linear motion stimuli, again comparable with results reported in previous studies (15,16).

We further tested whether the observed restriction of feature-based attentional modulation to only one feature dimension in MSTd is also present for spatial attention. The spatial attentional indices for spiral motion stimuli across all 105 neurons showed a significant spatial attentional modulation of 27% (p < 0.0001, signed-rank test). Similarly, we observed a significant spatial attentional modulation of 27% (p < 0.0001, signed-rank test) and 28% (p < 0.0001, signed-rank test) for spiral and linear motion stimuli respectively for the subset of 48 neurons that show multiplexed tuning, i.e. tuning to both spiral and linear motion patterns. This is in line with previous observations for spatial attentional effects in MST (7,36,37).

This specificity of feature-based attentional modulation in MST for only spiral motion stimuli, together with the spatial attention modulation of all stimuli supports our hypothesis that *feature-based* attention is only affecting those neuronal population that make a direct contribution to perception, while *spatial* attention acts as ‘passthrough’ gating mechanism dependent only upon receptive field location, affecting all stimuli within its spatial scope.

To evaluate the hypothesis that attentional modulation is absent for neuronal signals that are not directly linked to perception we assessed the link between the activity of neurons tuned to linear motion and behavioral reaction times. Correlations of neuronal firing rates with behavioral reaction time (i.e. shorter reaction times for trials with higher firing rates) have been proposed as evidence of a task-related recruitment of neurons in various region of visual cortex such as MT & VIP (38), LIP (39), FEF(40). These studies have demonstrated an inverse relationship of neuronal firing rates and reaction times. But so far these effects have been shown only for spatial attention paradigms. If feature-based attentional modulation is limited to those stimulus features to whose perception a given neuron contributes, then the relationship between firing rate and reaction time might also be restricted to stimulus feature(s) and dimension(s) for which a neuron demonstrates feature-based attentional modulation. We hypothesized that the distribution of firing rates indices (see Methods) would be skewed to positive values for spiral motion in MSTd and for linear motion in MT but not for linear motion in MSTd. This is because positive indices represent higher firing rates for short RT trials. 46 out of 48 neurons from area MSTd were used in the analysis for spiral motion and linear motion as two neurons had to be excluded due to insufficient number of trials with short RTs.

Responses to linear motion in MT were available for 60 neurons. The mean firing rate indeces for spiral and linear motion in MSTd and linear motion in MT (figure 3) are 0.016, −0.019 and 0.025 respectively. Significant shifts of firing rate indices to positive values were observed only for linear motion (in MT (p < 0.0001, figure 3) and spiral motion in MSTd (p = 0.0054, figure 3). The distribution is not significantly shifted to positive values for linear motion in MSTd (p = 0.9867, figure 3). This provides evidence for a contribution of area MST to the perception of spiral motion and area MT for the perception of linear motion. These results further support our hypothesis that feature-based attentional modulation only affects neurons which contribute to the perception of the given visual feature.

**Figure 3:**
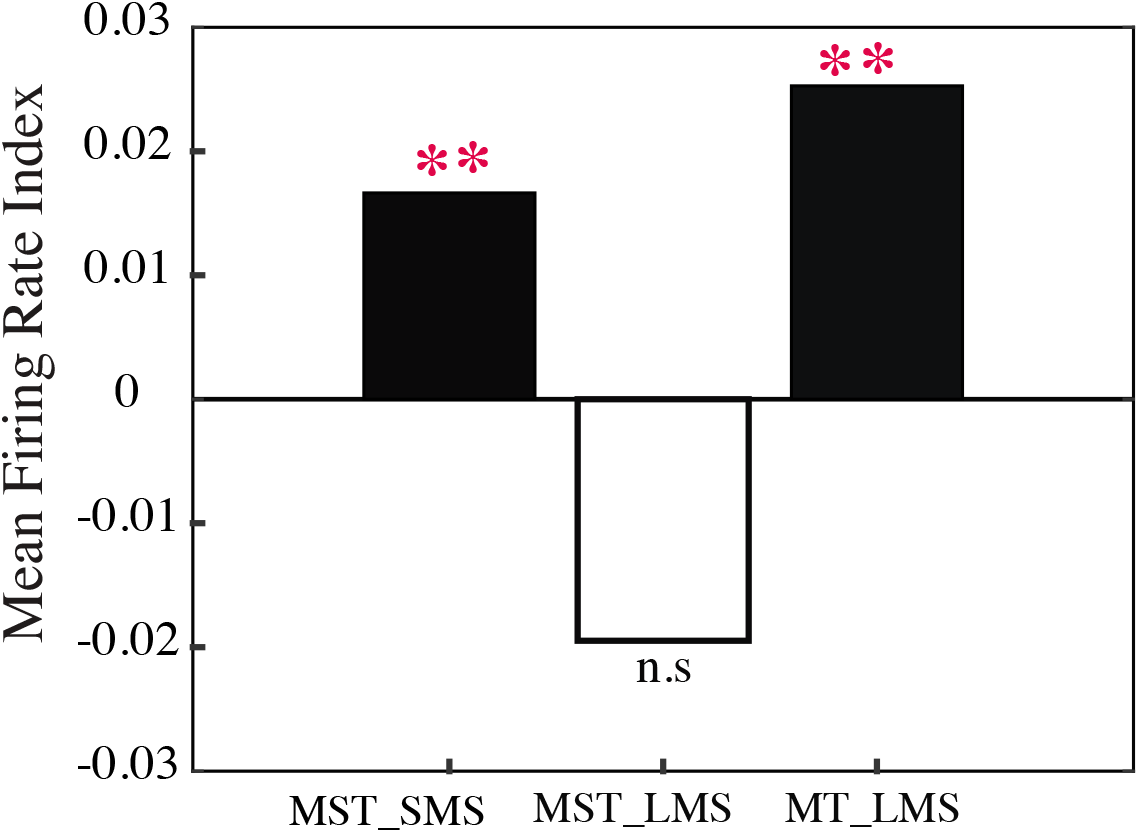
Firing rate index. Bar plot of the mean firing rate indices. Firing rate indices were calculated by dividing the difference of the firing rate for long RT trials from that of short RT trials by the sum of the two firing rates. Black bars represent the significantly positive mean indices for spiral motion stimuli in MSTd (0.0166, p = 0.0054) and linear motion in MT (0.0253, p < 0.0001). The white bar represents the mean firing rate index for linear motion in MSTd, which did not significantly deviate from zero (−0.019, p = 0.9867).

In summary, MSTd neurons with ‘multiplexed’ selectivity show tuning and spatial attentional modulation to both stimulus types (linear and spiral motion stimuli), but show feature-based attentional modulation for only spiral motion stimuli, These results support the hypothesis that neuronal tuning for a particular stimulus feature is not a sufficient criterion for a neuronal population’s causal contribution to this feature’s perception, even though the population might contribute to the overall representation of the stimulus.

## Discussion

Understanding the evolution of attentional modulation along the hierarchy of cortical visual information processing is crucial for an understanding of the interaction of bottom-up and top-down influences in sensory cortex. With this aim in mind we used the ‘multiplexed’, ‘tangled’ or ‘mixed’ (41–47) selectivity of many neurons in area MST of rhesus monkeys to spiral and linear visual motion to compare the modulation caused by spatial and feature-based attention for these two motion types.

Our data show a striking asymmetry between the feature-based attentional modulation of linear and spiral motion responses. While spatial attention affects both stimulus types with the same gain change to neuronal responses, feature-based attentional modulation in MST is only affecting the neuronal responses to spiral, but not linear motion stimuli, despite similar and robust tuning for these two stimulus dimensions. This asymmetry is complemented by a difference in the behavioral relevance of the two selectivities. Namely, we show that MST responses to spiral motion stimuli are linked to the animals’ behavioral performance (Fig. 3), while for linear motion responses we did not find evidence for such a link.

Our data thus demonstrate a dissociation between sensory preferences and attentional modulation in that they show for the first time that feature-based attentional modulation does not affect *all* stimulus parameters a cell in extrastriate cortex is tuned for. While the feature-similarity model of attention (15–17) in its current form correctly considers a similarity between a cell’s selectivity and the attended feature a *necessary* condition for a feature-based attentional enhancement it needs to be refined to account for our observation that such a similarity is not a *sufficient* condition. Rather, beyond the similarity it is necessary that a cell selective for an attended feature also directly contribute to the perception (and thus behavioral performance) of the feature.

This strong link of feature-based neuronal modulation with perception is well in line with recent observations from area MT in rhesus monkeys, suggesting that visual attention in primate visual cortex contributes to an enhanced internal representation of attended visual stimuli that is optimized for perceptual and behavioral performance, even at the expense of compromising an accurate stimulus representation (48,49).

The lack of a systematic feature-based attentional modulation of responses to linear motion stimuli we observed is particularly surprising, since we show (in the same animals, performing the same tasks), that such modulation is present in area MT, the extrastriate area providing the majority of the sensory input to MST. This demonstrates that attentional modulation can be lost along the hierarchical processing of information in visual cortex. We consider this evidence that the linear motion selectivity observed in MST is not ‘inherited’ from area MT but is generated ‘de novo’ in MST as part of the computation performed in MST to generate the complex spiral motion selectivity that we and others observed (25,50–53). While there is the general assumption that information available in one area of a visual cortical pathway is passed on downstream to the next area (confirmed by Movshon & Newsome (54) for the V1 to MT projection), this is not always the case. For example, Khawaja & Pack (55) showed that the selectivity for stationary plaids is not passed on from area MT to MST. A ‘de novo’ computation has been previously proposed for MST (27), where MST linear motion tuning is generated by complex weighing mechanisms of the input received from area MT neurons tuned to different directions.

The observation that MST linear motion responses are not directly linked to the perception of stimuli of this type seems to be in disagreement with a microstimulation study (36). In that study effects of microstimulation on linear motion perception were observed in area MSTd, using a paradigm that has been used successfully to directly link the responses of MT neurons to linear motion perception in rhesus monkeys (56). But note, that the paradigm used focuses on *spatial* attention. In agreement, our study (7,57) showed *spatial* attentional modulation for both linear and spiral motion in area MSTd.

These studies, as well as our data, thus support the view that spatial attention serves as a high-gain pass-through gating mechanism, optimizing the processing of sensory information from relevant locations in the environment (3, 51–54). In essence, spatial attention ensures that the full capabilities of sensory information processing are available to stimuli at attended locations, while the gain of neurons processing stimuli at unattended locations is actively reduced (14). This is in line with the proposal that spatial attention has a selection (i.e. gating) rather than a capacity-matching effect (3, 55, 56).

One might wonder, if our observation reflects a general inability of feature-based attention to modulate more than one feature at a time for a given neuron, possibly because of anatomical, pharmacological or network limitations of the specificity of top-down control of feature-based attentional modulation for mixed-selectivity. This concern is unwarranted, based on a study by Ruff and Born (63). They exploited the dual selectivity of individual MT neurons for binocular disparity and linear visual motion direction and observed feature-based attentional modulation for both features within the same neurons. The magnitude of feature-based attentional modulation was directly correlated to the tuning strength for binocular disparity (which was less than that for the linear motion stimuli). They concluded, that ‘feature attention effects can be found in neurons for multiple features including those to which they are not most strongly tuned for’.

In summary, our data support the notion of a highly optimized attentional system where the interplay of spatial and feature-based attention builds a location-based gating mechanism, providing a high-gain unimpeded pass-through of information from attended locations, combined with a feature-based modulation that differentially enhances the gain of only those neurons that are selective for the currently attended feature-set *and* contribute to its perception.

## Materials and Methods

Research with non-human primates represents a small but indispensable component of neuroscience research. The scientists in this study are aware and are committed to the great responsibility they have in ensuring the best possible science with the least possible harm to the animals (64). All animal procedures of this study were conducted in accordance with the applicable laws and regulations governing animal research and care and have been approved by the responsible regional government office (Niedersaechsisches Landesamt fuer Verbraucherschutz und Lebensmittelsicherheit (LAVES)) under the permit number 33.14.42502-04-064/07.

The two animals (*macaca mulatta*, male) were group-housed with other macaque monkeys in facilities of the German Primate Center in Goettingen, Germany in accordance with all applicable German and European regulations. The facility provides the animals with an enriched environment (incl. a multitude of toys and wooden structures (65,66), natural as well as artificial light), exceeding the size requirements of the European regulations, including access to outdoor space. Surgeries were performed aseptically under gas anesthesia using standard techniques, including appropriate peri-surgical analgesia and monitoring to minimize potential suffering. The German Primate Center has several staff veterinarians that regularly monitor and examine the animals and consult on procedures. During the study the animals had unrestricted access to food and fluid, except on the days where data were collected or the animals were trained on the behavioral paradigm. On these days the animals were allowed unlimited access to fluid through their performance in the behavioral paradigm. Here the animals received fluid rewards for every correctly performed trial. Throughout the study the animals’ psychological and veterinary welfare was monitored by the veterinarians, the animal facility staff and the lab’s scientists, all specialized in working with non-human primates.

We have established a comprehensive set of measures to ensure that the severity of our experimental procedures falls into the category of mild to moderate, according to the severity categorization of Annex VIII of the European Union’s directive 2010/63/EU on the protection of animals used for scientific purposes (see also (67)).

The two animals (male Macaca mulatta) participating in this study were healthy at the conclusion of our study and were subsequently used in other studies.

For the current study the animals performed a visual target detection task during daily training or recording sessions while seated in custom-made primate chairs at a viewing distance of 57cm to a CRT computer monitor presenting stimuli of 40° x 30° visual angle at a resolution of 40 pixel/deg. The animals reported subtle changes of relevant visual stimuli (luminance decrease of the fixation-dot during receptive field mapping and speed acceleration within random dot motion stimuli during the attention tasks) for liquid reward. Eye movements were recorded using video-based eye tracking (ET49, Thomas Recording, Giessen, Germany).

Following initial training, custom-made orthopedic implants were surgically placed onto the monkey’s skull preventing head movements during training and extracellular recording sessions. A recording chamber was placed on top of a surgically placed craniotomy over the left (monkey N: 3 mm posterior, 16 mm lateral; Crist Instruments, CILUX Recording Chamber 35°, Hagerstown, MD) or the right (monkey W: 3 mm posterior, 16 mm lateral; custom-fit computer-aided milled magnetic resonance imaging (MRI) compatible chamber, via digitized monkey skull surface reconstruction, 3di, Jena, Germany) parietal lobe. The location of the craniotomy was based on registration of the stereotaxic implant coordinates to that of animal’s pre-surgical MR images. Post-surgical MRIs verified the correct positioning and precise targeting of the area MST and MT via functional characterization based on expected white/gray matter transitions.

For extracellular recordings, up to three micro-electrodes were lowered with micrometer precision to the target area (Mini-Matrix, Thomas Recordings, Giessen, Germany). The raw electrode signal was amplified and filtered with a multi-channel processor (Map System, Plexon, Inc.), using headstages with unit gain. Spiking activity was obtained following a 100-8000 Hz bandpass filter and further amplification and digitization with a 40 kHz sampling rate. Spikes were isolated using a voltage-time window and all data analyzed in the current study came from well-isolated neurons. The local field potentials recorded are being analyzed in a separate study (69).

The experimental stimuli were moving random dot patterns (RDPs), which consisted of small bright dots (density: 8 dots per degree, luminance 75 cd/m^2^) plotted within a stationary circular aperture on a gray background of 35 cd/m^2^. Movement of the dots was created by the appropriate displacement of each dot at the monitor refresh rate of 75Hz. The RDPs stimuli were either linear motion stimuli (LMS) or spiral motion space (SMS) patterns. In linear motion stimuli the dots moved along parallel linear trajectories. Spiral motion stimuli come from a spiral motion space where expansion, clockwise rotation, contraction and counterclockwise rotation are neighboring stimuli, with a continuum of stimuli in between these cardinal directions (21). The direction of a specific SMS stimulus is determined by the angle that the direction of all of its individual dots form with radial reference lines. By varying this angle by equal steps we created the 12 directions within the SMS used in this study.

Recording of attention-related MT and MSTd single unit activity, was always preceded by the identification and characterization (speed and direction tuning profile) of the neurons receptive field (figure 1, represented by yellow dotted circle). Animals engaged in a luminance task on a centrally positioned fixation point, while an unattended RDP was presented within the location of the receptive filed of the neuron under study. The size of the RDP was matched to allow the placement of two RDPs at equal eccentricity to the fixation point (inside & outside the receptive field). To determine direction tuning for SMS, twelve motion directions (in steps of 30°) with a velocity of 8 degrees per second for the dots furthest away from the center were randomly chosen in intervals of 827ms. For linear motion stimuli eight directions (in steps of 45 degrees) were used.

Single unit responses to each direction were defined as a mean firing rate in an interval of 80-800ms after onset of a particular motion direction. Both linear and spiral direction tuning curves were calculated online and fitted with a circular Gaussian across the mean responses to the twelve stimuli directions. The motion direction yielding the highest mean firing rate was then presented at eight different speeds (between 0.5 and 64 deg/sec) to determine the preferred speed of each neuron. Parts of our data were also analyzed for effects of burstiness in the firing patterns (68).

After mapping of the receptive field, and determining the preferred direction and speed, the animals’ task was switched to the attentional paradigm. In brief, the task (Figure 1a) required the animals to foveate a static central visual stimulus (fixation point − 0.2° x 0.2°), maintain their gaze upon this target throughout the trial. After foveating the fixation point, a visual cue flashed briefly (67ms) at the peripheral target location followed by 400ms of a blank screen (used to determine a neuron’s baseline firing-rate). Next, two incoherent RDPs were presented at equal eccentricity on either side of the fixation point for 375ms. One stimulus was always positioned inside the receptive field (RF) of the neuron under study, the other in the opposite visual hemifield. Then, two coherent motion stimuli (RDP) were presented for 250-2500ms at exact same locations where incoherent motion RDPs were presented before. The size, speed and direction of motion of the RDP were optimized to match the preferences of the neuron for coherent motion either in spiral (SMS) (25) or linear motion space (LM). At the beginning of each trial one of the two stimulus locations was cued using a stationary stimulus (cue), indicating the upcoming target. The animals’ task was to correctly detect a brief speed increase in the target stimulus at a random point in time while ignoring changes of the other stimulus (the distractor). Trials ended when the animal gave a response and received a liquid reward for each correct detection. When no response to the target change was given within a response time window or when the animal diverted its gaze from the fixation point no reward was given. In a given trial the stimulus inside and outside the receptive field were of the same type, i.e. either spiral or linear motion stimuli (fig 1C and 1D). Neuronal responses in two attentional conditions were recorded, manipulating either the state of feature-based or of spatial attention.

For the feature-based attentional modulation, spatial attention was held constant, that is, in all trials the target was the RDP outside the receptive field (moving in either the preferred or the anti-preferred direction), with the distractor inside the receptive field always moving in the preferred direction. This paradigm allows comparing responses when feature-based attention was directed at the preferred vs. the anti-preferred direction outside the receptive field, without changing the sensory stimulus inside the receptive field. This paradigm has proven to be very suitable for determining feature-based attentional modulation (15,24,41,48,69). The time course of single unit responses for the two feature-based attentional conditions is shown in figure 1C and 1D for spiral and linear motion stimuli respectively.

For the spatial attentional modulation, feature-based attention was held constant, with the target and distractor always moving in the preferred direction of the neuron. The effect of spatial attention was determined by comparing neuronal responses when the stimulus inside the receptive field was the target and when it was the distractor, i.e. when attention was directed into or out of the receptive field. The time course of responses of a single unit for the two attentional conditions is shown in figure 1B for spiral motion stimuli.

Neural activities were recorded from area MT and MSTd. From area MSTd a total of 105 neurons were recorded for the spatial and feature-based attention paradigms with SMS. Out of these 105 neurons, for a sub-set of 48 neurons, spatial and feature-based attention paradigms were recorded for LMS also. From area MT a total of 60 neurons were recorded for the spatial and feature-based attention paradigms with LMS only.

Data were analyzed offline with custom scripts using MATLAB (The Math Works, Natick, MA). For the analysis of neuronal data only correctly performed, completed trials were included. To estimate spike rates, spike density functions (SDF) were evaluated for all correctly performed, complete trials of different attentional conditions, by convolving each spike in a trial with a Gaussian function (σ = 30). The SDFs of trials from the same attentional conditions were averaged and the firing rates were evaluated by taking the mean of the averaged SDFs over a time window of 600 ms starting from 300 ms after the onset of coherent motion stimuli, corresponding to the period of sustained activity (Gray shaded area in figure 2A & 2C).

Attentional modulation was evaluated using an attentional index, i.e. by dividing the difference in firing rate between two attentional conditions by their sum. Spatial attention modulation was measured by dividing the difference of the firing rates when attention was directed to preferred direction inside and outside the receptive field, to their sum. While, feature-based attention was evaluated by dividing the difference of firing rates when attention was directed to the preferred and the anti-preferred direction outside the receptive field to their sum.

It has been shown in various area of visual cortex that, the firing rate of a neuron is inversely related to the reaction time (RT) of an animal (38–40,70). This link has been interpreted as a direct contribution of an individual neuron to the organism’s perception. We investigated the presence of such a relationship between firing rate and reaction times in our data set, in an epoch of 300ms before the response event (speed increment in target stimuli) to which monkeys were trained to respond. This analysis was performed on trials where the preferred direction (SMS/LMS) was presented both inside and outside the receptive field, while attention was directed outside the receptive field. First, we divided the correctly completed trials of a given neuron into those with RTs shorter and longer than the median RT. Only those trials were included in the analysis where the response event happened not earlier than 400ms after the coherent motion onset. This was done to ensure that firing rates were sampled from periods of sustained firing. Next, for each of the two trial types (long and short RTs), firing rates were sampled from SDFs for an epoch of 300ms before the response event, with a sliding window of 30ms starting with the response event and shifting in steps of 15ms, giving 20 bins of firing rates for each trial. The firing rate in each of the 20 bins corresponding to long and short RTs were then averaged across trials for each neuron. Firing rates indices were then computed for each of the 20 bins by dividing the difference between the firing rates for long RT trials from those of short RT trials by the sum of the two firing rates. These data were then pooled across all neurons.

All data presented here and the associated analysis codes will be made available in a public repository when this manuscript is published as a peer-reviewed publication

## Acknowledgements

This project was supported by grants to ST from the Deutsche Forschungsgemeinschaft (DFG): Research Unit 1847 (project A1) “Physiology of Distributed Computing Underlying Higher Brain Functions in Non-Human Primates” and Collaborative Research Center 889 (project C4) “Cellular Mechanisms of Sensory Processing” as well as a grant (01GQ0433) of the German Federal Ministry of Education and Research to the Bernstein Center for Computational Neuroscience, Goettingen. We are thankful for expert technical, administrative and animal care support in particular from Ralf Brockhausen, Kevin Windolph, Dirk Pruesse, Leonore Burchardt, Sina Pluemer, Beatrix Glaser and Janine Kramer.

## References

1. Treue S. Visual attention: the where, what, how and why of saliency. Curr Opin Neurobiol. 2003;13(4):428–32.

2. Anton-Erxleben K, Stephan VM, Treue S. Attention reshapes center-surround receptive field structure in macaque cortical area MT. Cereb Cortex. 2009;19(10):2466–78.

3. Chen Y, Seidemann E. Attentional modulations related to spatial gating but not to allocation of limited resources in primate V1. Neuron. 2012;74(3):557–66.

4. Niebergall R, Khayat PS, Treue S, Martinez-Trujillo JC. Expansion of MT neurons excitatory receptive fields during covert attentive tracking. J Neurosci. 2011;31(43):15499–510.

5. Posner MI, Snyder CR, Davidson BJ. Attention and the detection of signals. J Exp Psychol Gen. 1980;109(2):160.

6. Spitzer H, Desimone R, Moran J. Increased attention enhances both behavioral and neuronal performance. Science (80-). 1988;240(4850):338–40.

7. Treue S, Maunsell JHR. Attentional modulation of visual motion processing in cortical areas MT and MST. Nature. 1996;382(6591).

8. Womelsdorf T, Anton-Erxleben K, Pieper F, Treue S. Dynamic shifts of visual receptive fields in cortical area MT by spatial attention. Nat Neurosci. 2006;9(9):1156–60.

9. Womelsdorf T, Anton-Erxleben K, Treue S. Receptive field shift and shrinkage in macaque middle temporal area through attentional gain modulation. J Neurosci. 2008;28(36):8934–44.

10. Esghaei M, Daliri MR, Treue S. Attention decreases phase-amplitude coupling, enhancing stimulus discriminability in cortical area MT. Front Neural Circuits. 2015;9:82.

11. Khamechian MB, Kozyrev V, Treue S, Esghaei M, Daliri MR. Routing information flow by separate neural synchrony frequencies allows for “functionally labeled lines” in higher primate cortex. Proc Natl Acad Sci. 2019;116(25):12506–15.

12. McAdams CJ, Maunsell JHR. Effects of attention on orientation-tuning functions of single neurons in macaque cortical area V4. J Neurosci. 1999;19(1):431–41.

13. Williford T, Maunsell JHR. Effects of spatial attention on contrast response functions in macaque area V4. J Neurophysiol. 2006;96(1):40–54.

14. Malek N, Treue S, Khayat P, Martinez-Trujillo J. Distracter suppression dominates attentional modulation of responses to multiple stimuli inside the receptive fields of middle temporal neurons. Eur J Neurosci. 2017;46(12):2844–58.

15. Treue S, Trujillo JCM. Feature-based attention influences motion processing gain in macaque visual cortex. Nature. 1999;399(6736):575.

16. Katzner S, Busse L, Treue S. Attention to the color of a moving stimulus modulates motion-signal processing in macaque area MT: evidence for a unified attentional system. Front Syst Neurosci. 2009;3:12.

17. Katzner S, Busse L, Treue S. Feature-based attentional integration of color and visual motion. J Vis. 2006;6(3):7.

18. Liu T, Stevens ST, Carrasco M. Comparing the time course and efficacy of spatial and feature-based attention. Vision Res. 2007;47(1):108–13.

19. Ling S, Liu T, Carrasco M. How spatial and feature-based attention affect the gain and tuning of population responses. Vision Res. 2009;49(10):1194–204.

20. Ibbotson MR, Crowder NA, Cloherty SL, Price NSC, Mustari MJ. Saccadic modulation of neural responses: possible roles in saccadic suppression, enhancement, and time compression. J Neurosci. 2008;28(43):10952–60.

21. Ibos G, Freedman DJ. Interaction between spatial and feature attention in posterior parietal cortex. Neuron. 2016;91(4):931–43.

22. Yao T, Treue S, Krishna BS. Saccade-synchronized rapid attention shifts in macaque visual cortical area MT. Nat Commun. 2018;9(1):958.

23. Yao T, Ketkar M, Treue S, Krishna BS. Visual attention is available at a task-relevant location rapidly after a saccade. Elife. 2016;5:e18009.

24. Yao T, Treue S, Krishna BS. An attention-sensitive memory trace in macaque MT following saccadic eye movements. PLoS Biol. 2016;14(2):e1002390.

25. Graziano MS, Andersen RA, Snowden RJ. Tuning of MST neurons to spiral motions. J Neurosci. 1994;14(1):54–67.

26. Celebrini S, Newsome WT. Neuronal and psychophysical sensitivity to motion signals in extrastriate area MST of the macaque monkey. J Neurosci. 1994;14(7):4109–24.

27. Lappe M, Bremmer F, Pekel M, Thiele A, Hoffmann K-P. Optic flow processing in monkey STS: a theoretical and experimental approach. J Neurosci. 1996;16(19):6265–85.

28. Wild B, Treue S. Primate extrastriate cortical area MST: a gateway between sensation and cognition. J Neurophysiol. 2021;125(5):1851–82.

29. Dubin MJ, Duffy CJ. Behavioral influences on cortical neuronal responses to optic flow. Cereb Cortex. 2006;17(7):1722–32.

30. Recanzone GH, Wurtz RH. Effects of attention on MT and MST neuronal activity during pursuit initiation. J Neurophysiol. 2000;83(2):777–90.

31. Britten KH, Shadlen MN, Newsome WT, Movshon JA. The analysis of visual motion: a comparison of neuronal and psychophysical performance. J Neurosci. 1992;12(12):4745–65.

32. Logothetis NK, Schall JD. Neuronal correlates of subjective visual perception. Science (80-). 1989;245(4919):761–3.

33. Polonsky A, Blake R, Braun J, Heeger DJ. Neuronal activity in human primary visual cortex correlates with perception during binocular rivalry. Nat Neurosci. 2000;3(11):1153.

34. Martinez-Trujillo JC, Treue S. Feature-based attention increases the selectivity of population responses in primate visual cortex. Curr Biol. 2004;14(9):744–51.

35. McAdams CJ, Maunsell JHR. Attention to both space and feature modulates neuronal responses in macaque area V4. J Neurophysiol. 2000;83(3):1751–5.

36. Celebrini S, Newsome WT. Microstimulation of extrastriate area MST influences performance on a direction discrimination task. J Neurophysiol. 1995;73(2):437–48.

37. O’Craven KM, Rosen BR, Kwong KK, Treisman A, Savoy RL. Voluntary attention modulates fMRI activity in human MT–MST. Neuron. 1997;18(4):591–8.

38. Cook EP, Maunsell JHR. Dynamics of neuronal responses in macaque MT and VIP during motion detection. Nat Neurosci. 2002;5(10):985–94.

39. Janssen P, Shadlen MN. A representation of the hazard rate of elapsed time in macaque area LIP. Nat Neurosci. 2005;8(2):234–41.

40. Everling S, Munoz DP. Neuronal correlates for preparatory set associated with pro-saccades and anti-saccades in the primate frontal eye field. J Neurosci. 2000;20(1):387–400.

41. Barak O, Rigotti M, Fusi S. The sparseness of mixed selectivity neurons controls the generalization–discrimination trade-off. J Neurosci. 2013;33(9):3844–56.

42. Rigotti M, Barak O, Warden MR, Wang X-J, Daw ND, Miller EK, et al. The importance of mixed selectivity in complex cognitive tasks. Nature. 2013;497(7451):585.

43. DiCarlo JJ, Zoccolan D, Rust NC. How does the brain solve visual object recognition? Neuron. 2012;73(3):415–34.

44. Rust NC, DiCarlo JJ. Selectivity and tolerance (“invariance”) both increase as visual information propagates from cortical area V4 to IT. J Neurosci. 2010;30(39):12978–95.

45. Sereno AB, Maunsell JHR. Shape selectivity in primate lateral intraparietal cortex. Nature. 1998;395(6701):500.

46. Pagan M, Urban LS, Wohl MP, Rust NC. Signals in inferotemporal and perirhinal cortex suggest an untangling of visual target information. Nat Neurosci. 2013;16(8):1132.

47. Parthasarathy A, Herikstad R, Bong JH, Medina FS, Libedinsky C, Yen S-C. Mixed selectivity morphs population codes in prefrontal cortex. Nat Neurosci. 2017;20(12):1770–9.

48. Kozyrev V, Daliri MR, Schwedhelm P, Treue S. Strategic deployment of feature-based attentional gain in primate visual cortex. PLoS Biol. 2019;17(8).

49. Mehrpour V, Martinez-Trujillo JC, Treue S. Attention amplifies neural representations of changes in sensory input at the expense of perceptual accuracy. Nat Commun. 2020;11(1):1–8.

50. Beardsley SA, Ward RL, Vaina LM. A neural network model of spiral–planar motion tuning in MSTd. Vision Res. 2003;43(5):577–95.

51. Grossberg S, Mingolla E, Pack C. A neural model of motion processing and visual navigation by cortical area MST. Cereb Cortex. 1999;9(8):878–95.

52. Mineault PJ, Khawaja FA, Butts DA, Pack CC. Hierarchical processing of complex motion along the primate dorsal visual pathway. Proc Natl Acad Sci. 2012;109(16):E972–80.

53. Saito H, Yukie M, Tanaka K, Hikosaka K, Fukada Y, Iwai E. Integration of direction signals of image motion in the superior temporal sulcus of the macaque monkey. J Neurosci. 1986;6(1):145–57.

54. Movshon JA, Newsome WT. Visual response properties of striate cortical neurons projecting to area MT in macaque monkeys. J Neurosci. 1996;16(23):7733–41.

55. Khawaja FA, Liu LD, Pack CC. Responses of MST neurons to plaid stimuli. J Neurophysiol. 2013;110(1):63–74.

56. Salzman CD, Britten KH, Newsome WT. Cortical microstimulation influences perceptual judgements of motion direction. Nature. 1990;346(6280):174.

57. Treue S, Maunsell JHR. Effects of attention on the processing of motion in macaque middle temporal and medial superior temporal visual cortical areas. J Neurosci. 1999;19(17):7591–602.

58. Vidyasagar TR. Gating of neuronal responses in macaque primary visual cortex by an attentional spotlight. Neuroreport. 1998;9(9):1947–52.

59. Moore T, Armstrong KM. Selective gating of visual signals by microstimulation of frontal cortex. Nature. 2003;421(6921):370.

60. Wang X-J, Yang GR. A disinhibitory circuit motif and flexible information routing in the brain. Curr Opin Neurobiol. 2018;49:75–83.

61. Grothe I, Rotermund D, Neitzel SD, Mandon S, Ernst UA, Kreiter AK, et al. Attention selectively gates afferent signal transmission to area V4. J Neurosci. 2018;38(14):3441–52.

62. Palmigiano A, Geisel T, Wolf F, Battaglia D. Flexible information routing by transient synchrony. Nat Neurosci. 2017;20(7):1014.

63. Ruff DA, Born RT. Feature attention for binocular disparity in primate area MT depends on tuning strength. J Neurophysiol. 2015;113(5):1545–55.

64. Roelfsema PR, Treue S. Basic neuroscience research with nonhuman primates: a small but indispensable component of biomedical research. Neuron. 2014;82(6):1200–4.

65. Berger M, Calapai A, Stephan V, Niessing M, Burchardt L, Gail A, et al. Standardized automated training of rhesus monkeys for neuroscience research in their housing environment. J Neurophysiol. 2017;119(3):796–807.

66. Calapai A, Berger M, Niessing M, Heisig K, Brockhausen R, Treue S, et al. A cage-based training, cognitive testing and enrichment system optimized for rhesus macaques in neuroscience research. Behav Res Methods. 2017;49(1):35–45.

67. Pfefferle D, Plümer S, Burchardt L, Treue S, Gail A. Assessment of stress responses in rhesus macaques (Macaca mulatta) to daily routine procedures in system neuroscience based on salivary cortisol concentrations. PLoS One. 2018;13(1):e0190190.

68. Xue C, Kaping D, Ray SB, Krishna BS, Treue S. Spatial attention reduces burstiness in macaque visual cortical area MST. Cereb Cortex. 2017;27(1):83–91.

69. Patzwahl DR, Treue S. Combining spatial and feature-based attention within the receptive field of MT neurons. Vision Res. 2009;49(10):1188–93.

70. Womelsdorf T, Fries P, Mitra PP, Desimone R. Gamma-band synchronization in visual cortex predicts speed of change detection. Nature. 2006;439(7077):733–6.

69 Aboutorabi E, Ray SB, Kaping D, Shahbazi F, Treue S, Esghaei E. Attentional shift of individual neurons’ activity relative to the neighbouring population dynamics explains attentional improvement of behaviour. 14^th^ Goettingen Meeting of German Neuroscience Society. 2021

