## Supplementary Table for "Feature-based gating of cortical information transmission"

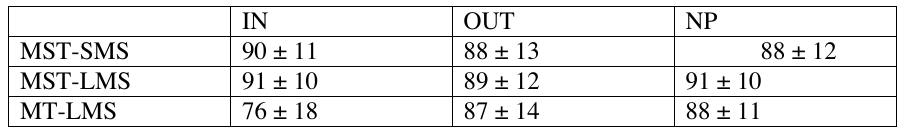

**Behavioral Performance**

Average behavioral performance of the two monkeys in different conditions that were used to evaluate spatial and feature based attentional indices. IN, corresponds to condition where attention was directed to preferred direction inside the RF, while preferred direction was also presented outside the RF; OUT corresponds to condition where again preferred direction was presented both inside and outside the RF but attention was directed to outside the RF; NP corresponds to condition where attention was directed to anti-preferred direction outside the RF while preferred direction was presented inside the RF. MST-SMS represented data of 105 neurons recorded form area MSTd with SMS; MST-LMS represents data from 48 neurons recorded form area MSTd with LMS; MT-LMS corresponds to data from 60 neurons recorded from area MT with LMS. The value in each cell represents performance in percentage (mean ± SD).
