## Supplementary figures and images for "Feature-based gating of cortical information transmission"

### Supplementary Figure 1

**Supplementary figure 1**

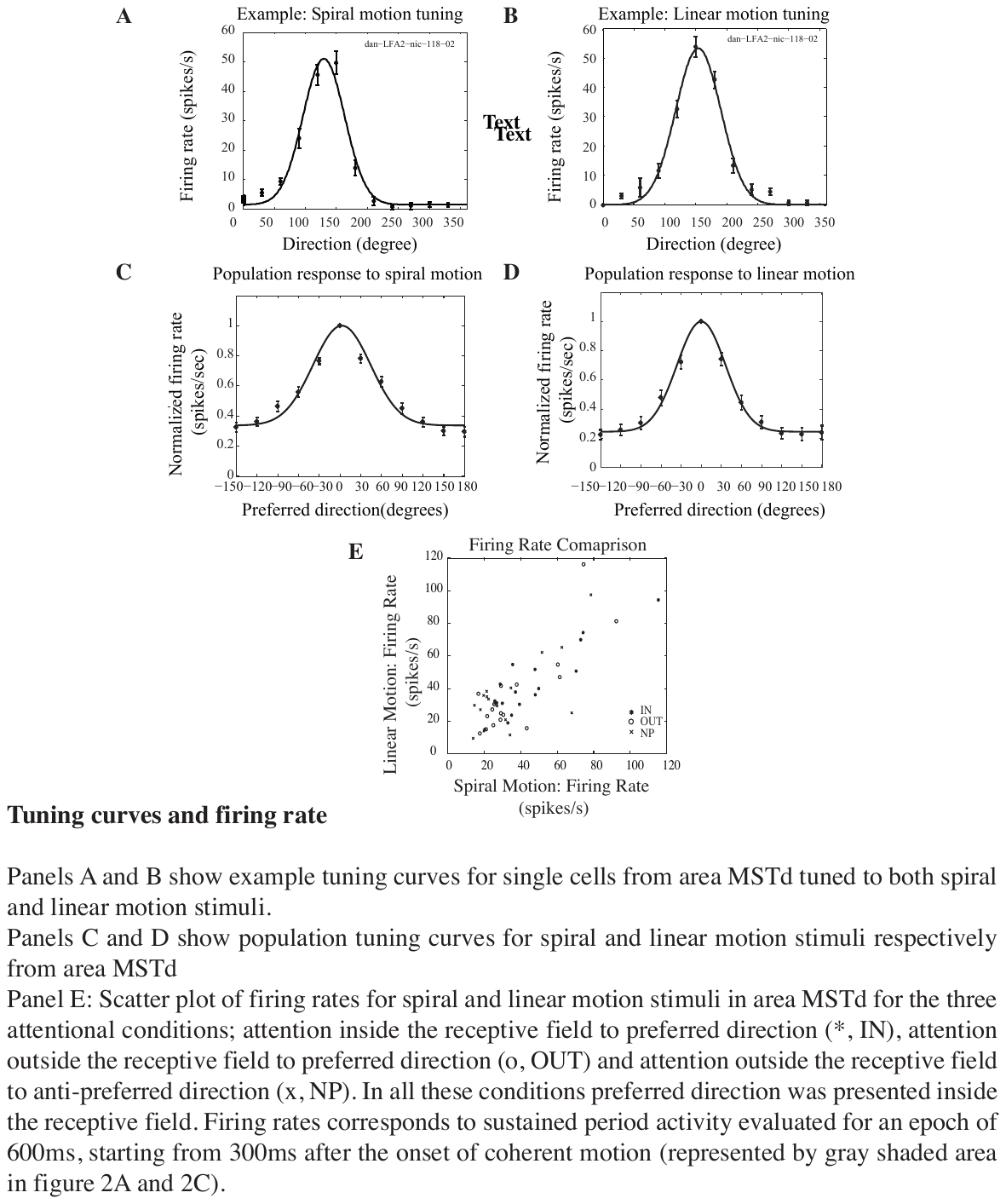
