## Supplementary Figure 2 for "Feature-based gating of cortical information transmission"

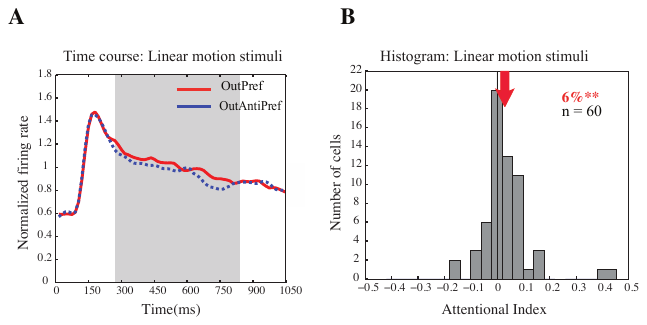

**Feature-based attentional modulation in area MT.**

**A.** Normalized spike density function of a population of 60 neurons from area MT, for two conditions recorded to measure feature-based attention for linear motion stimuli: Attention outside the receptive field to preferred direction (red curve) and attention outside the receptive field to anti-preferred direction (blue dotted). Grey shaded area represents 600ms time period of sustained activity starting after 300ms of onset. This time period was chosen to evaluate attentional indices for the units recorded.

**B.** Histogram showing distribution of attentional indices for feature-based attention of 60 neurons recorded from area MT with linear motion stimuli. Mean attentional index was 0.030 (represented by red arrow) corresponding to a significant modulation of 6% (Sign-rank test, p<0.001).
